# The prolyl hydroxylase PHD3 maintains β-cell glucose metabolism during fatty acid excess

**DOI:** 10.1101/2020.04.30.068106

**Authors:** Daniela Nasteska, Federica Cuozzo, Alpesh Thakker, Rula Bany Bakar, Rebecca Westbrook, Ildem Akerman, James Cantley, Daniel A. Tennant, David J. Hodson

## Abstract

The alpha ketoglutarate-dependent dioxygenase, prolyl-4-hydroxylase 3 (PHD3), is a hypoxia-inducible factor target that uses molecular oxygen to hydroxylate proline. While PHD3 has been reported to influence cancer cell metabolism and liver insulin sensitivity, relatively little is known about effects of this highly conserved enzyme in insulin-secreting β-cells. Here, we show that deletion of PHD3 specifically in β-cells (βPHD3KO) is associated with impaired glucose homeostasis in mice fed high fat diet. In the early stages of dietary fat excess, βPHD3KO islets energetically rewire, leading to defects in the management of pyruvate fate and a shift away from glycolysis. However, βPHD3KO islets are able to maintain oxidative phosphorylation and insulin secretion by increasing utilization of fatty acids to supply the tricarboxylic acid cycle. This nutrient-sensing switch cannot be sustained and βPHD3KO islets begin to show signs of failure in response to prolonged metabolic stress, including impaired glucose-stimulated ATP/ADP rises, Ca^2+^ fluxes and insulin secretion. Thus, PHD3 might be a pivotal component of the β-cell glucose metabolism machinery by suppressing the use of fatty acids as a primary fuel source, under obesogenic and insulin resistant states.

**SIGNIFICANCE STATEMENT:** Prolyl-4-hydroxylase 3 (PHD3) is involved in the oxygen-dependent regulation of cell phenotype. A number of recent studies have shown that PHD3 might operate at the interface between oxygen availability and metabolism. To understand how PHD3 influences insulin secretion, which depends on intact glucose metabolism, we generated mice lacking PHD3 specifically in pancreatic β-cells. These mice, termed βPHD3KO, are apparently normal until fed high fat diet at which point their β-cells switch to fatty acids as a fuel source. This switch cannot be tolerated and β-cells in βPHD3KO mice eventually fail. Thus, PHD3 maintains glucose-stimulated insulin secretion in β-cells during states of fatty acid excess, such as diabetes and obesity.

## INTRODUCTION

The prolyl-hydroxylase domain proteins (PHD1-3) encoded for by the *Egl-9* homologue (*EGLN*) genes are alpha ketoglutarate-dependent dioxygenases, which regulate cell function by catalyzing hydroxylation of prolyl residues within various substrates using molecular oxygen (1-4). There are three well-described mammalian isozymes: PHD1, PHD2 and PHD3, which were originally described as hydroxylating the alpha subunit of the transcription factor Hypoxia-Inducible Factor (HIF) under normoxia (4), thus targeting it for polyubiquitylation and proteasomal degradation. When oxygen concentration becomes limited, PHD activity decreases and HIF is stabilized, leading to dimerization with the beta subunit and transcriptional regulation of target genes regulating the cellular response to hypoxia (5). While PHDs are generally regarded to be master HIF regulators, it is becoming increasingly apparent that they target a range of other substrates influencing cell function (6-9).

PHD3 is unusual amongst the PHDs: it is transcriptionally regulated by HIF1 during hypoxia (10) although it does not always act to destabilize HIF1 (11, 12). A number of roles for PHD3 have been described under conditions of stress or hypoxia, including: macrophage influx and neutrophil survival (13, 14), apoptosis in various cancer models (8, 15, 16), and tumor cell survival (9) (reviewed in (17)). Due to the dependence of PHD3 on alpha-ketoglutarate and oxygen for its activity (18), many of these actions are likely to be mediated through alterations in cell metabolism (19). Indeed, PHD3 increases glucose uptake in cancer cells through interactions with pyruvate kinase M2 (8, 20). In tumors exhibiting mutations in succinate dehydrogenase, fumarate hydratase and isocitrate dehydrogenase 1 and 2 (21-23), PHD3 activity is altered by aberrantly high cytosolic concentrations of succinate, fumarate and 2-hydroxyglutarate (2-HG), suggesting that inactivation of this enzyme might be involved in the cellular transformation process. PHD3 has more recently been shown to hydroxylate and activate acetyl-CoA carboxylase 2 (ACC2), defined as the fatty acid oxidation gatekeeper, thus decreasing fatty acid breakdown and restraining myeloid cell proliferation during nutrient abundance (24). Together, these studies place PHD3 as a central player in the regulation of glucose and fatty acid utilization with clear implications for metabolic disease risk.

Along these lines, PHD3 has been reported to influence insulin sensitivity in the liver (25, 26), as well as maintain glucose-stimulated insulin secretion in a rat β-cell line (27). However, little is known about how PHD3 might contribute to glucose homeostasis and diabetes risk through effects directly in primary pancreatic β-cells. To ensure the appropriate release of insulin, β-cells have become well-adapted as glucose sensors. Thus, glucose enters the β-cell by facilitated diffusion through low affinity glucose transporters (28), before conversion into glucose-6-phosphate by glucokinase and subsequent splitting into pyruvate (29). The pyruvate then undergoes oxidative metabolism in the mitochondrial matrix through the tricarboxylic acid (TCA) cycle, driving increases in ATP/ADP ratio and leading to closure of ATP-sensitive K^+^ channels (30). This cascade triggers membrane depolarization, opening of voltage-dependent Ca^2+^ channels, influx of Ca^2+^, and Ca^2+^-dependent exocytosis of insulin vesicles through interactions with the SNARE machinery (30). Together with repression of hexokinase, monocarboxylic acid transporter 1 and lactate dehydrogenase A (31, 32), stimulus-secretion coupling prevents the inappropriate release of insulin in response to low glucose, amino acids or lactate.

Given its reported roles in dictating fuel preference, we hypothesized that PHD3 might function as a pivotal component of the β-cell glucose-sensing machinery by suppressing the use of fatty acids as an energy source (27). To further investigate PHD3-regulated β-cell function in depth, we subjected a model of β-cell-specific *Egln3*, encoding for PHD3, deletion to extensive *in vivo* and *in vitro* characterization, including detailed stable isotope-resolved metabolic tracing. Here, we show that loss of PHD3 causes metabolic remodelling in the early stages of metabolic stress by shifting β-cell fuel source from glucose to fatty acids. However, this metabolic switch is overwhelmed as fatty acids accumulate, ultimately leading to β-cell failure. As such, PHD3 is likely to constitute a fundamental mechanism to restrain fatty acid utilization and maintain glucose-sensing in β-cells during early stages of metabolic stress and insulin resistance.

## RESULTS

### PHD3 knockout does not induce a hypoxic gene expression phenotype

We first generated a model of β-cell PHD3 knockout (βPHD3KO) by crossing the *Ins1Cre* deleter strain (33) with animals harboring flox’d alleles for *Egln3* (34), which encodes PHD3. Recombination efficiency of the *Ins1Cre* line was verified in-house by crossing to mTmG reporter animals and was found to be >90%. Gene expression analyses showed a 2-fold reduction in *Egln3* in βPHD3KO islets (Figure 1A). Western blotting revealed a similar ∼50% knockdown of PHD3 protein in βPHD3KO islets (Figure 1B), the remainder most likely reflecting the relatively higher levels of *Egln3* detected in α-cells, as shown by RNA-seq (35, 36). While we attempted immunofluorescence staining, we could not detect a specific signal in β-cells, probably reflecting known sensitivity issues with PHD3 antibodies. Previous studies have shown that PHD3 is highly regulated at the transcriptomic level by hypoxia (10), and in line with this, we also found that *Egln3* levels in hypoxic (1% O_2_) βPHD3CON islets were highly upregulated (Figure 1C). While *Egln3* is expressed at low abundance in sorted β-cells (35, 36), this is likely to be a result of profound re-oxygenation following dissociation, thus suppressing *Egln3* expression (37).

**Figure 1:**
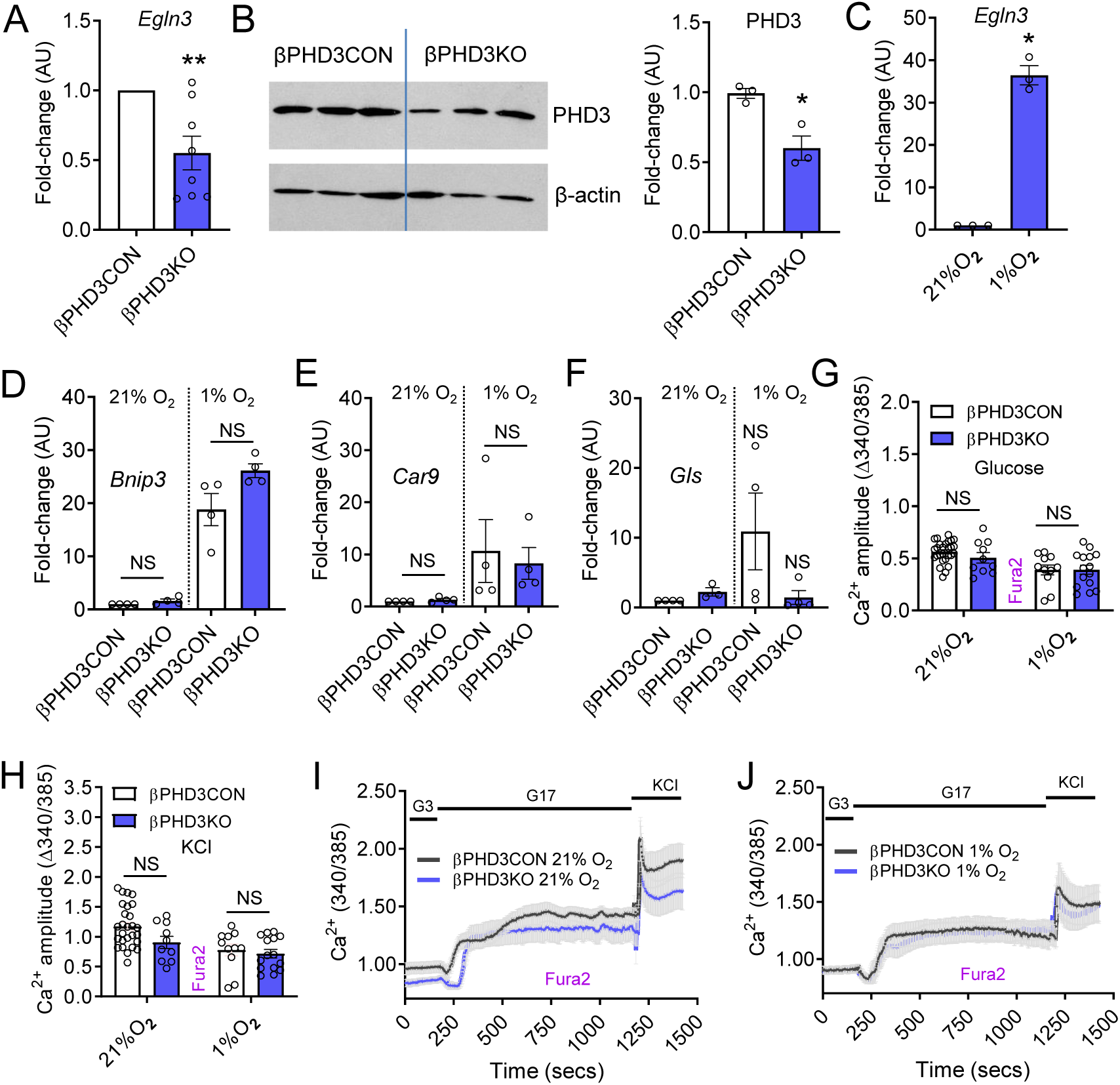
Generation and validation of mice lacking PHD3 in pancreatic β-cells. (A) *Egln3* expression is reduced in islets of βPHD3KO mice versus control (βPHD3CON) littermates (n = 8 animals) (paired t-test). (B) Western blot analyses showing decreased PHD3 expression in βPHD3KO mice (n = 100-150 islets from 3 animals run on the same gel) (paired t-test). (C) *Egln3* expression is highly upregulated following exposure of islets to hypoxic (1% O_2_) conditions for 24 hrs (n = 3 animals) (paired t-test). (D-F) Expression of the HIF1α-target genes *Bnip* (D), *Car9* (E) or *Gls* (F), is similar or decreased in βPHD3KO versus βPHD3CON islets exposed to normoxia (21% O_2_) or hypoxia (1% O_2_) 24 hrs (n = 4 animals) (Kruskal-Wallis test Dunn’s multiple comparison test) (G and H) Glucose- (G) and KCl- (H) stimulated Ca^2+^ fluxes are not significantly different in βPHD3KO versus βPHD3CON islets exposed to normoxia or hypoxia (n = 11-27 islets, 2 animals/genotype) (two-way ANOVA; Sidak’s multiple comparison test). (I and J) Mean Ca^2+^ traces from βPHD3CON (I) and βPHD3KO (J) islets exposed to normoxia or hypoxia. Bar graphs show scatter plot with mean ± SEM. Line graphs show mean ± SEM. *P<0.05, **P<0.01 and NS, non-significant. PHD3, prolyl-hydroxylase 3.

To account for HIF-dependent effects on β-cell phenotype in βPHD3KO animals, a number of canonical HIF1α-target genes were assessed. Notably, levels of *Bnip3* and *Car9* were similar between normoxic (21% O_2_) βPHD3CON and βPHD3KO islets (Figure 1D-F). Further suggesting the presence of intact HIF signaling, *Bnip3, Car9* were upregulated to similar levels in hypoxic (1% O_2_) βPHD3CON and βPHD3KO islets (Figure 1D-F), while Gls did not reliably increase (Figure 1D-F). Lastly, glucose and KCl-stimulated Ca^2+^ fluxes, shown to be sensitive to HIF stabilization (38), were similar in βPHD3CON and βPHD3KO islets exposed to hypoxia (Figure 1G-J).

### PHD3 does not contribute to glucose homeostasis and insulin release under normal diet

Male and female βPHD3KO mice presented with normal growth curves from 8-18 weeks of age compared to βPHD3CON littermates (Figure 2A and B). Glucose tolerance testing in the same animals showed no abnormalities in glycemia (Figure 2C and D), which did not change up until the age of 20 weeks (Figure 2E and F). As expected, βPHD3KO mice displayed similar insulin sensitivity (Figure 2G), islet size distribution and β-cell mass (Figure 2H-J) to their βPHD3CON littermates.

**Figure 2:**
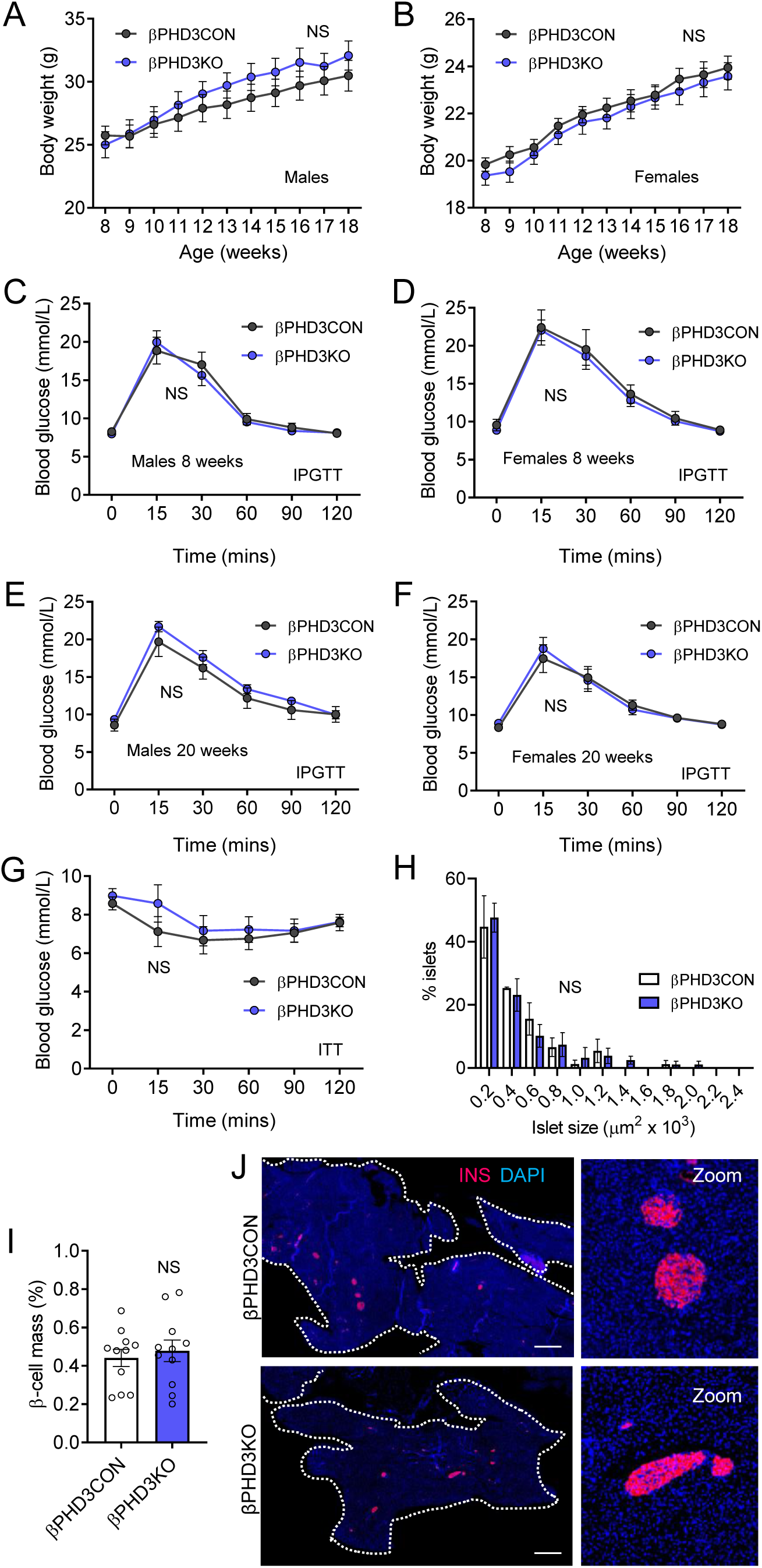
βPHD3KO *in vivo* phenotype under standard chow conditions. (A and B) Male (A) and female (B) βPHD3CON and βPHD3KO mice possess similar growth curves and adult body weight (n = 8-10 male and 15 female animals/genotype) (two-way repeated measures ANOVA; Sidak’s multiple comparison test). (C and D) No differences in intraperitoneal glucose tolerance are detected between βPHD3CON and βPHD3KO male (n = 13 animals/genotype) (C) and female (n = 10 animals/genotype) (D) 8-week-old mice (two-way repeated measures ANOVA; Sidak’s multiple comparison test). (E and F) No differences in intraperitoneal glucose tolerance are detected between βPHD3CON and βPHD3KO male (E) and female (F) 20-week-old mice (two-way repeated measures ANOVA; Sidak’s multiple comparison test) (n = 8-16 male and 8 female animals/genotype). (G) Insulin sensitivity is similar in βPHD3CON and βPHD3KO mice (n = 5-6 animals/genotype) (two-way repeated measures ANOVA; Sidak’s multiple comparison test). (H-J) Cell resolution reconstruction of entire pancreatic sections shows no differences in islet size and β-cell mass between βPHD3CON and βPHD3KO mice. Quantification is shown in (H and I), with representative images in (J) (scale bar = 530 µm) (inset is a zoom showing maintenance of cellular resolution in a single image) (n = 3 animals/genotype) (unpaired t-test). Bar graphs show scatter plot with mean ± SEM. Line graphs show mean ± SEM. *P<0.05, **P<0.01 and NS, non-significant. PHD3, prolyl-hydroxylase 3.

Isolation of islets for more detailed *in vitro* workup revealed normal expression of the β-cell transcription factors/differentiation markers *Pdx1, Mafa* and *Nkx6-1* in βPHD3KO islets (Figure 3A-C). Further suggestive of mature β-cell function, live imaging approaches revealed intact glucose-stimulated ATP/ADP ratios (Figure 3D and E) and Ca^2+^ fluxes (Figure 3F and G) in βPHD3KO islets. While glucose-stimulated insulin secretion was similar in islets isolated from βPHD3CON and βPHD3KO animals, responses to the incretin-mimetic Exendin-4 (Ex4) were blunted (Figure 3H). This defect in Ex4-potentiated insulin secretion was not due to reductions in *Glp1r* expression (Figure 3I) or cAMP responses to the incretin-mimetic (Figure 3J and K). Moreover, oral glucose tolerance, largely determined by incretin release from the intestine (39), was similar in βPHD3CON and βPHD3KO mice (Figure 3L).

**Figure 3:**
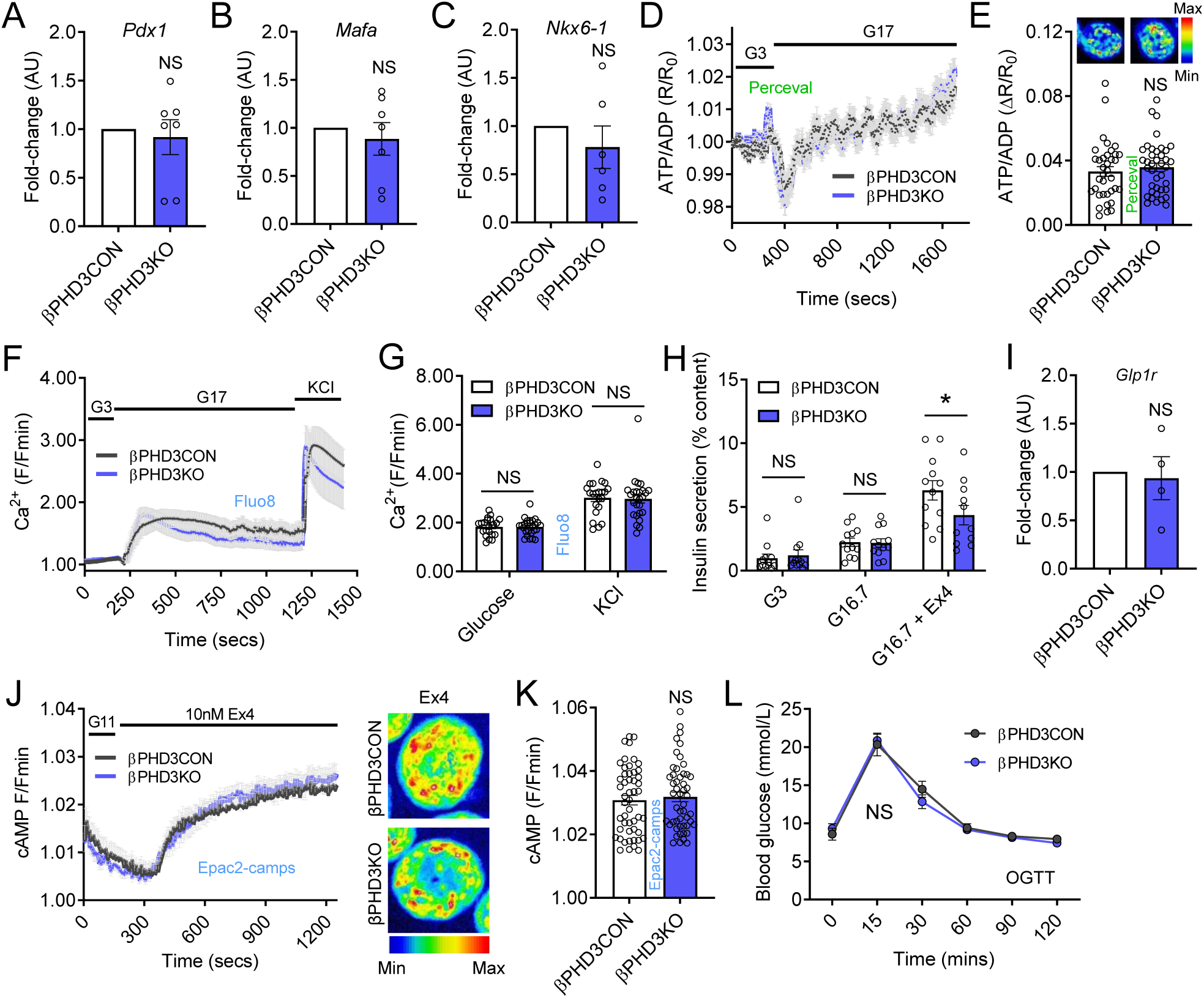
βPHD3KO *in vitro* phenotype under standard chow conditions. (A-C) Expression of the β-cell maturity markers *Pdx1* (A), *Mafa* (B) and *Nkx6-1* (C) is similar in βPHD3CON and βPHD3KO islets (n = 6-7 animals) (paired t-test). (D and E) Glucose-stimulated ATP/ADP rises do not differ in islets of βPHD3CON and βPHD3KO mice, as shown by mean traces (D) and summary bar graph (E) (representative images shown above each bar; a single islet has been cropped for clarity) (n = 38-40 islets, 4-5 animals/genotype) (unpaired t-test). (F and G) Glucose- and KCl-stimulated Ca^2+^ rises do not differ in islets of βPHD3CON and βPHD3KO mice, as shown by mean traces (F), as well as summary bar graph (G) (n = 22-26 islets, 2 animals/genotype) (two-way ANOVA; Sidak’s multiple comparison test). (H) Insulin secretory responses to Exendin-4 (Ex4), but not glucose, are impaired in βPHD3KO islets (n = 12 replicates from 3 animals/genotype) (two-way ANOVA; Bonferonni’s multiple comparison test). (I) *Glp1r* expression is similar in βPHD3CON and βPHD3KO islets (n = 4 animals/genotype) (paired t-test). (J and K) cAMP responses to Ex4 do not differ between βPHD3CON and βPHD3KO islets, as shown by mean traces and summary bar graph (representative images shown; a single islet has been cropped for clarity) (n = 50 islets from 4-5 animals/genotype) (unpaired t-test). (L) Oral glucose tolerance is almost identical in βPHD3CON and βPHD3KO mice (n = 4 animals/genotype) (two-way repeated measures ANOVA; Sidak’s multiple comparison test). Bar graphs show scatter plot with mean ± SEM. Line graphs show mean ± SEM. *P<0.05, **P<0.01 and NS, non-significant. Exendin-4, Ex4; PHD3, prolyl-hydroxylase 3; G3, 3 mM glucose; G11, 11 mM glucose; G16.7, 16.7 mM glucose; G17, 17 mM glucose.

### Loss of PHD3 improves insulin secretion at the onset of metabolic stress

We next examined whether PHD3 might play a more important role in regulating insulin release during metabolic stress. Indeed, the increase in islet size that occurs during insulin resistance is associated with a hypoxic state (40), expected to increase PHD3 levels via HIF1 activity (41). Therefore, animals were placed on high fat diet (HFD) to induce obesity and metabolic stress(42).

Following 4 weeks HFD, *Egln3* was mildly but significantly upregulated in βPHD3CON islets (Figure 4A). As expected, *Egln3* levels remained suppressed in 4 weeks HFD βPHD3KO islets (Figure 4A). Glucose tolerance testing revealed significantly impaired glucose homestasis in βPHD3KO mice at 4 weeks but not at 72 hours HFD (Figure 4B and C), despite similar body weight gain compared to βPHD3CON littermates (Figure 4D). As expected, fasting blood glucose levels were elevated in βPHD3CON mice following 4 weeks HFD (Figure 4E). There was no effect of Cre or flox’d alleles *per se* on glucose tolerance following 4 weeks HFD (Figure 4F). Glucose-stimulated insulin secretion *in vivo* was however increased in 4 weeks HFD βPHD3KO mice (Figure 4G), suggesting that glucose intolerance might be associated with hyperinsulinemia (43). These increases in circulating insulin were associated with an almost 2-fold increase in β-cell mass in 4 weeks HFD βPHD3KO mice (Figure 4H and I), associated with a significant increase in the proportion of larger islets (Figure 4J). In addition, islets isolated from the same animals secreted significantly more insulin in glucose-stimulated and Ex4-potentiated states (Figure 4K). Increased islet size or insulin expression induced by HFD were unlikely to account for the overall increase in *in vitro* insulin secretion, since all measures were corrected for insulin content. *Bnip3* and *Gls* levels remained unchanged (Figure 4L and M), while *Car9* was downregulated (Figure 4N), suggesting that HIF1α stabilization was unlikely to be a major feature in HFD βPHD3KO islet.

**Figure 4:**
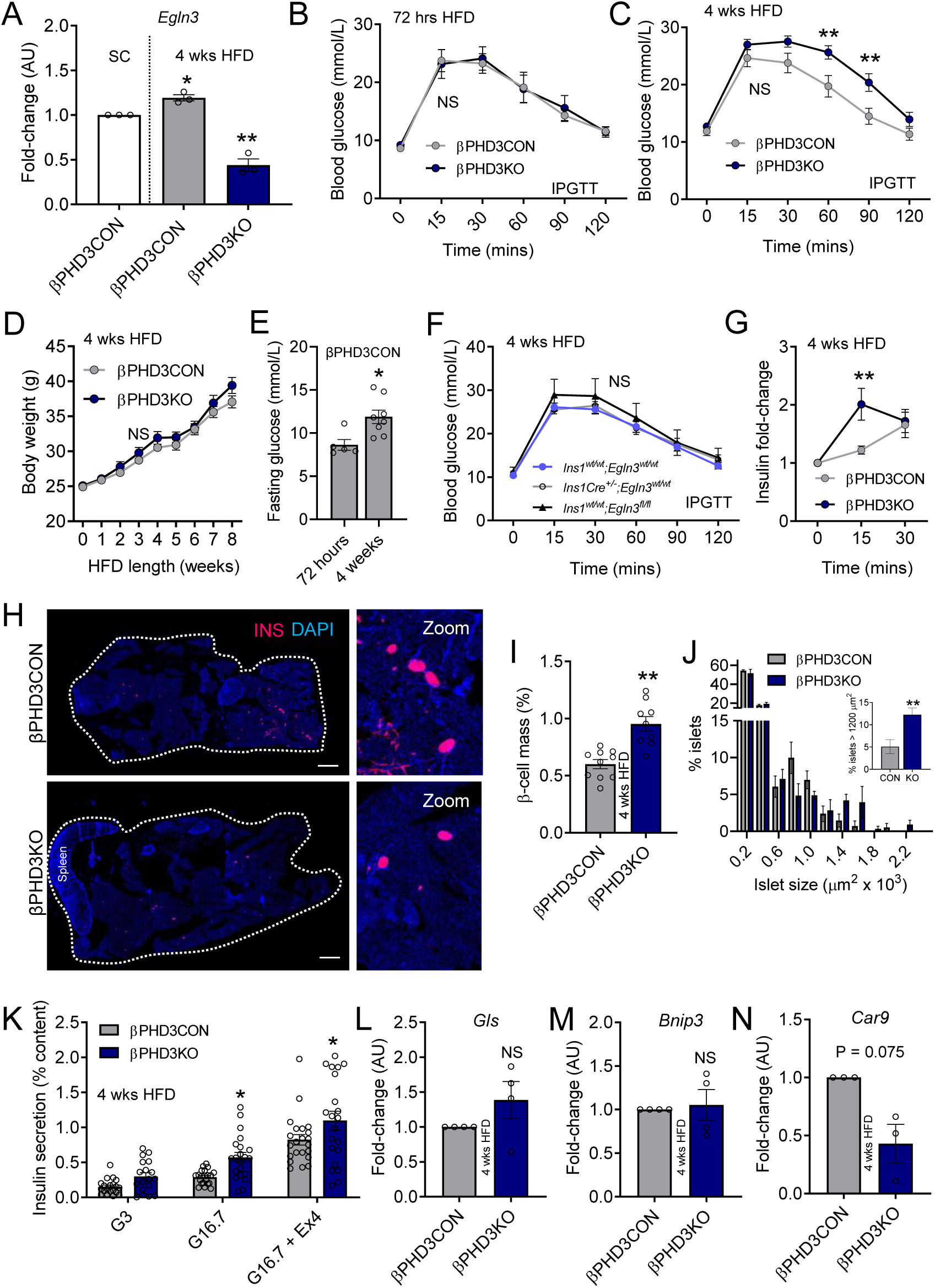
βPHD3KO *in vivo* and *in vitro* phenotype during metabolic stress. (A) *Egln3* expression is upregulated in 4 weeks HFD βPHD3CON, but not βPHD3KO islets (n = 3 animals/genotype) (one-way ANOVA; Sidak’s multiple comparison test). (B and C) Glucose tolerance in male βPHD3KO mice is unaffected by 72 hrs HFD (n = 5 animals/genotype), but is markedly impaired by 4 weeks HFD (C) (n = 8-11 animals/genotype) (two-way repeated measures ANOVA; Sidak’s multiple comparison test). (D) Growth curves and adult body weight are similar in male βPHD3CON and βPHD3KO animals fed HFD (n = 11-12 animals/genotype) (two-way repeated measures ANOVA; Sidak’s multiple comparison test). (E) Fasting blood gucose levels are significantly higher in male βPHD3CON after 4 weeks versus 72 hrs HFD (n = 5-8 animals) (unpaired t-test). (F) Glucose tolerance is unaffected in male *Cre*- and *Egln3*^fl/fl^-only controls (n = 10-13 animals/genotype) (two-way repeated measures ANOVA; Tukey’s multiple comparison test). (G) Insulin secretory responses to intraperitoneal glucose are higher in fasted male βPHD3KO mice versus βPHD3CON littermates fed HFD for 4 weeks (n = 6-13 animals/genotype) (two-way repeated measures ANOVA; Sidak’s multiple comparison test). (H-J) Cell resolution reconstruction of entire pancreatic sections shows a 2-fold increase in β-cell mass, with a shift toward larger islets, in βPHD3KO compared to βPHD3CON mice following 4 weeks HFD. Representative images are shown in (H), with quantification in (I and J) (scale bar = 530 µm) (inset is a zoom showing maintenance of cellular resolution in a single image) (unpaired t-test) (n = 3 animals/genotype). (K) Glucose- and Exendin-4-stimulated insulin secretion is increased in islets from 4 weeks HFD βPHD3KO mice (n = 20 replicates from 4 animals/genotype) (two-way ANOVA; Bonferonni’s multiple comparison test). (L-N) Expression of the HIF1α-target genes *Gls* (L), *Bnip3* (M) and *Car9* (N) is either unchanged or decreased in 4 weeks HFD βPHD3KO islets (n = 3-4 animals/genotype) (paired t-test). Bar graphs show scatter plot with mean ± SEM. Line graphs show mean ± SEM. *P<0.05, **P<0.01 and NS, non-significant. PHD3, prolyl-hydroxylase 3; Exendin-4, Ex4; G3, 3 mM glucose; G16.7, 16.7 mM glucose.

Thus, βPHD3KO mice are glucose intolerant on HFD, but their islets are larger and show improved insulin secretion. These data raise the possibility that nutrient-sensing and utilization might be altered in βPHD3KO islets.

### PHD3 maintains glucose metabolism in β-cells

Given the reported roles of PHD3 in glycolysis, we wondered whether the changes in β-cell function observed during the early phases of high fat feeding in βPHD3KO mice might be associated with changes in glucose metabolism. We first looked at glycolytic fluxes using ^14^C glucose. While glucose oxidation was not altered at low or high glucose in islets from 4 weeks HFD βPHD3KO mice (Figure 5A), there was a small but significant decrease in ^14^C content in the aqueous phase, indicating a net reduction in tricarboxylic acid (TCA) cycle/other metabolites derived from glycolysis (Figure 5B). Notably, a 2-fold reduction in incorporation of glucose into the lipid pool (i.e. glucose-driven lipogenesis) was also detected in 4 weeks HFD βPHD3KO islets (Figure 5C), suggestive of decreased glycolytic flux through the TCA cycle and acetyl-CoA carboxylase 1 (ACC1) (44).

**Figure 5:**
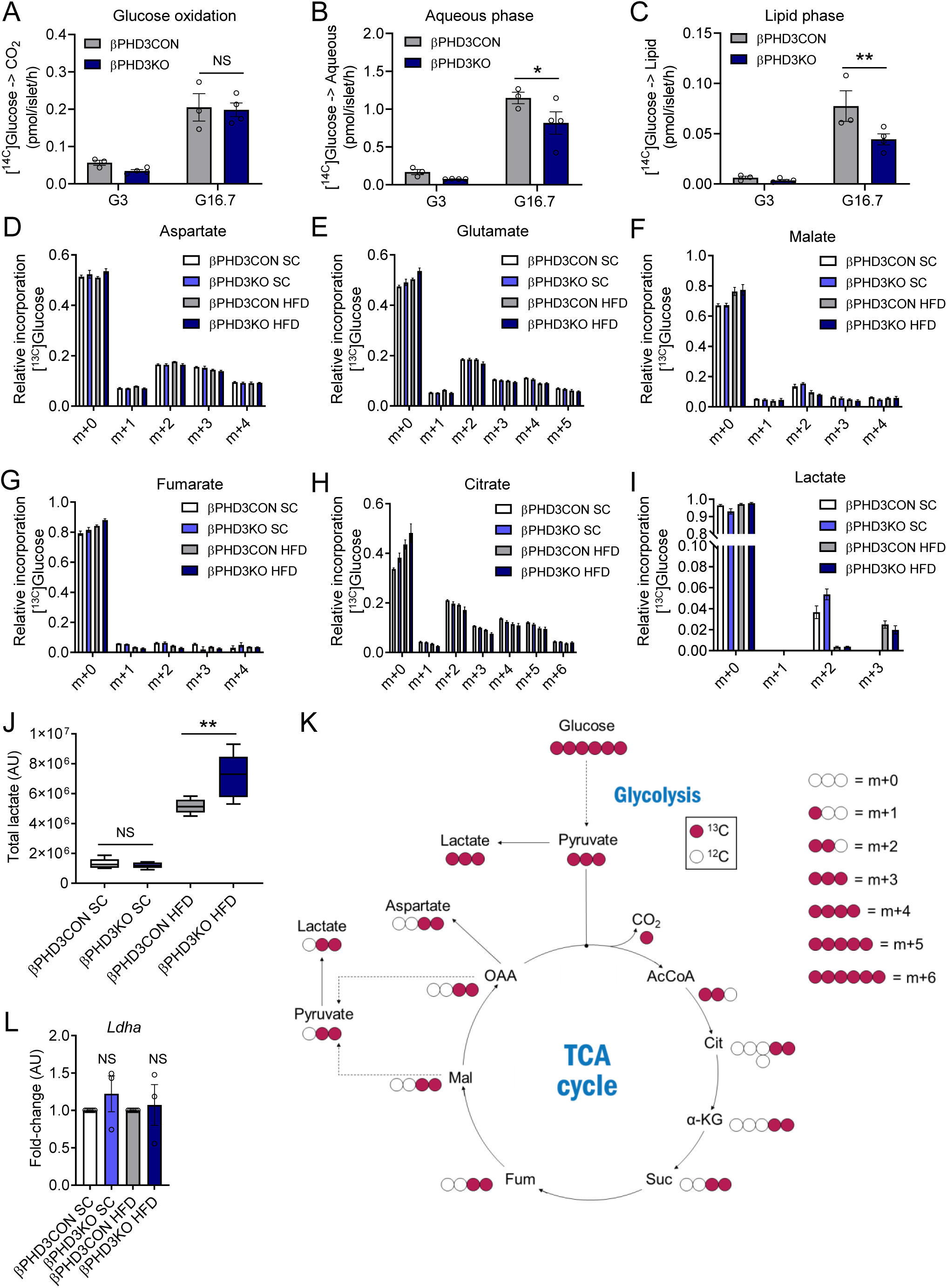
Metabolic rewiring in βPHD3KO islets during metabolic stress. (A-C) βPHD3KO islets possess intact glucose oxidation (A), but impaired accumulation of glycolytic/TCA cycle metabolites (B) and glucose-driven lipogenesis (C) following 4 weeks of HFD (n = 3 islet preparations from 3 animals/genotype) (two-way ANOVA, Fisher LSD). (D-H) Mass isotopomer distribution (MID) showing that ^13^C_6_ incorporation from glucose into aspartate (D), glutamate (E), malate (F), fumarate (G) or citrate (H) is similar in βPHD3CON and βPHD3KO islets isolated from mice fed SC or HFD (n = 3 replicates from 6 animals/genotype) (two-way ANOVA, Tukey’s multiple comparison test). (I) ^13^C_6_ glucose is incorporated primarily into m+2 lactate in SC βPHD3CON and βPHD3KO islets, whereas a shift to m+3 lactate is seen during 4 weeks HFD (n = 3 replicates from 3-6 animals/genotype) (two-way ANOVA, Tukey’s multiple comparison test). (J) Steady-state lactate levels are increased in βPHD3KO versus βPHD3CON islets following 4 weeks HFD (n = 3 animals/genotype) (one-way ANOVA, Bonferroni’s multiple comparison test). (K) Schematic showing fate of ^13^C_6_ glucose in βPHD3KO islets. (L) *Ldha* expression is similar in both SC and HFD βPHD3KO and βPHD3CON islets (n = 4 animals/genotype) (paired t-test). Bar graphs show mean ± SEM. *P<0.05, **P<0.01 and NS, non-significant. PHD3, prolyl-hydroxylase 3; *Ldha*; gene lactate dehydrogenase A.

To gain a higher resolution analysis of glucose fate, stable isotope-resolved tracing was performed in βPHD3KO islets using ^13^C_6_-glucose. GC-MS-based ^13^C_6_ mass isotopomer distribution analysis showed no differences in glucose incorporation into aspartate, glutamate, malate, fumarate or citrate in either standard chow or 4 weeks HFD βPHD3CON and βPHD3KO islets (Figure 5D-H). Thus, while the contribution of glucose to aqueous cellular metabolite pools is clearly reduced in 4 weeks HFD βPHD3KO islets (Figure 5B), there is no net change in the incorporation of glucose into each metabolite i.e. the TCA cycle proceeds normally despite lowered glucose fluxes. Islets from animals fed standard chow showed m+2 lactate accumulation (Figure 5I), which is consistent with lactate normally produced as a result of oxidative metabolism of glucose-derived pyruvate. However, during HFD there was a pronounced switch to reduction of pyruvate to lactate (indicated by the m+3 isotopomer) in both genotypes.

Further analysis of steady-state lactate levels showed a significant increase in lactate production in islets from HFD-fed βPHD3KO versus βPHD3CON mice (Figure 5J). Together with the m+2 → m+3 switch, this finding confirms initial measures with ^14^C glucose indicating reduced fueling of the TCA cycle by glycolysis (Figure 5K). Furthermore, the tracing data suggest that 4 weeks HFD βPHD3KO islets increase the reduction of pyruvate to support continued glycolysis through regeneration of the cytosolic NAD^+^ pool. The source of the lactate was unlikely to be through increases in expression of the “disallowed” gene lactate dehydrogenase A (*Ldha*) (31, 32), since *Ldha* levels were unchanged between βPHD3CON and βPHD3KO islets (Figure 5L).

Together, these data suggest that metabolic stress induces defects in the management of pyruvate fate in βPHD3KO islets, implying that insulin secretion must be maintained and even amplified through mechanisms other than glycolysis *in vitro*.

### PHD3 suppresses fatty acid use under metabolic stress

We hypothesized that βPHD3KO islets might switch to an alternative energy source to sustain their function, namely beta oxidation of fatty acids, which are present in excess during HFD. Moreover, in cancer cells PHD3 has been shown to increase activity of ACC2, which converts acetyl-CoA → malonyl-CoA, the latter suppressing carnitine palmitoyltransferase I (CPT1), the rate-limiting step in fatty acid oxidation (24). Indicating a profound change in β-cell nutrient preference, supplementation of culture medium with the fatty acid palmitate for 48 hours augmented glucose-stimulated and Ex4-potentiated insulin secretion in 4 weeks HFD βPHD3KO islets (Figure 6A). By contrast, 4 weeks HFD βPHD3CON islets showed no increase in glucose-stimulated insulin release following culture with palmitate (Figure 6B), confirming that the fatty acid was unlikely to induce lipotoxicity at the concentration and timing used here. Interestingly, 48 hrs incubation with palmitate increased Ex4-potentiated insulin secretion in 4 weeks HFD βPHD3CON islets (Figure 6B).

**Figure 6:**
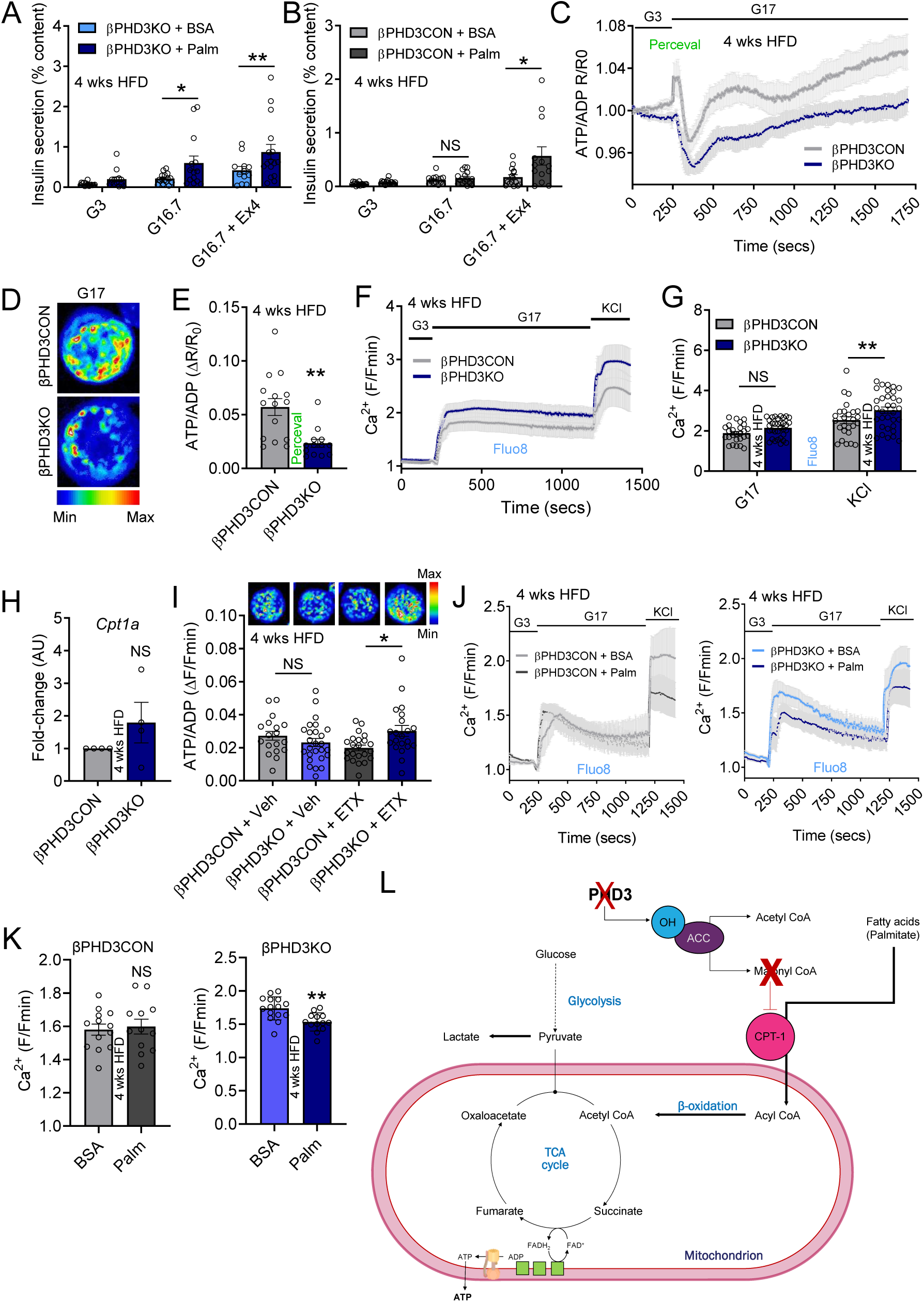
Nutrient preference is altered in βPHD3KO islets during metabolic stress. (A) Incubation of 4 weeks HFD βPHD3KO islets for 48 hrs with exogenous palmitate leads to enhanced glucose- and Exendin-4-stimulated insulin secretion (n = 12-17 replicates from 4 animals/genotype) (two-way ANOVA, Fisher LSD). (B) As for (A), but 4 weeks HFD βPHD3CON islets showing no changes in glucose-stimulated insulin secretion, but an increase in responses to Exendin-4 (n = 13-17 replicates from 4 animals/genotype) (two-way ANOVA, Fisher LSD). (C-E) Glucose-stimulated ATP/ADP rises are impaired in 4 weeks HFD βPHD3KO islets, as shown by mean traces (C), representative images (a single islet has been cropped for clarity) (D) and summary bar graph (E) (n = 13-15 islets from 4 animals/genotype) (unpaired t-test). (F and G) Glucose- and KCl-stimulated Ca^2+^ rises are similar (glucose) or increased (KCl) in βPHD3CON and βPHD3KO islets following 4 weeks HFD, as shown by mean traces (F) and summary bar graph (G) (n = 26-33 islets from 3 animals/genotype) (two-way measures ANOVA; Sidak’s multiple comparison test). (H) *Cpt1a* expression tends to be increased in 4 weeks HFD βPHD3KO islets (n = 4 animals/genotype) (paired t-test). (I) Inhibition of CPT1 using etomoxir (ETX) increases the amplitude of glucose-stimulated ATP/ADP ratio in 4 weeks HFD βPHD3KO islets (representative images shown above each bar; a single islet has been cropped for clarity) (n = 18 islets from 2 animals/genotype) (two-way measures ANOVA; Sidak’s multiple comparison test). (J and K) Ca^2+^ responses to glucose are impaired in 4 weeks HFD βPHD3KO islets, but not 4 weeks HFD βPHD3CON islets, following 48 hrs incubation with the free fatty acid palmitate (palm) (150 µM), as shown by mean traces (J) and summary bar graphs (K) (n = 12-15 islets from 2-3 animals/genotype) (unpaired t-test). (L) Schematic showing proposed effects of PHD3 deletion on β-cell metabolism under HFD. Bar graphs show scatter plot with mean ± SEM. Line graphs show mean ± SEM. *P<0.05, **P<0.01 and NS, non-significant. PHD3, prolyl-hydroxylase 3; *Cpt1a*, carnitine palmitoyltransferase 1A; Exendin-4, Ex4; G3, 3 mM glucose; G16.7, 16.7 mM glucose; G17, 17 mM glucose; BSA, bovine serum albumin; Palm, palmitate.

Further confirming a switch away from glycolysis, glucose-stimulated ATP/ADP ratios were markedly decreased in 4 weeks HFD βPHD3KO islets (Figure 6C-E), despite the increased insulin secretion (Figure 4K). While downstream Ca^2+^ fluxes were apparently normal in 4 weeks HFD βPHD3KO islets, this was likely due to increased sensitivity of voltage-dependent Ca^2+^ channel to membrane depolarization, since responses to KCl were significantly elevated (Figure 6F and G). Suggesting that CPT1 activity might be upregulated in 4 weeks HFD βPHD3KO islets, experiments were performed at high glucose, normally expected to inhibit CPT1 and fatty acid utilization through generation of malonyl-CoA, and mRNA levels of *Cpt1a* tended to be increased (Figure 6H). Moreover, application of the CPT1 inhibitor etomoxir was able to augment ATP/ADP responses to glucose in 4 weeks HFD βPHD3KO but not in βPHD3CON islets (Figure 6I). In line with this finding, culture with low palmitate concentrations decreased glucose-stimulated Ca^2+^ fluxes in 4 weeks HFD βPHD3KO (Figure 6J) but not in βPHD3CON islets (Figure 6K), presumably due to increased flux of acetyl-CoA into the TCA cycle.

Thus, following 4 weeks HFD, βPHD3KO islets become less reliant on glycolysis to fuel ATP/ADP production, are able to sustain oxidative phosphorylation through fatty acid oxidation, and secrete more insulin when both glucose and fatty acids are present. These changes, which are in agreement with our initial *in vivo* and *in vitro* phenotyping data (Figure 4), are shown schematically in Figure 6L.

### ACC1 and ACC2 are differentially regulated at the promoter level in β-cells

ACC1 and ACC2 are enzymes that catalyze the carboxylation of acetyl-CoA to malonyl-CoA, the rate-limiting step in fatty acid synthesis. β-cells are thought to predominantly express ACC1 (45, 46), which supplies cytosolic malonyl-CoA to fatty acid synthase for *de novo* lipid biosynthesis rather than for beta oxidation (44). By contrast, β-cells are reported to express negligible levels of ACC2, which inhibits CPT1 through generation of mitochondrial malonyl-CoA to suppress use of fatty acids via beta oxidation (45, 46).

To explore potential mechanisms that might underlie the changes occurring in 4 weeks HFD βPHD3KO islets, we decided to interrogate multiple published bulk islet and purified β-cell gene expression datasets (35, 47, 48). Notably, *ACACB* (encoding ACC2) was found to be present in β-cells, albeit at lower levels than *ACACA* (encoding ACC1) (Figure 7A). The expression levels of *ACACB* were significant, however, reaching similar levels to the β-cell transcription factor *HNF1A* (Figure 7A).

**Figure 7:**
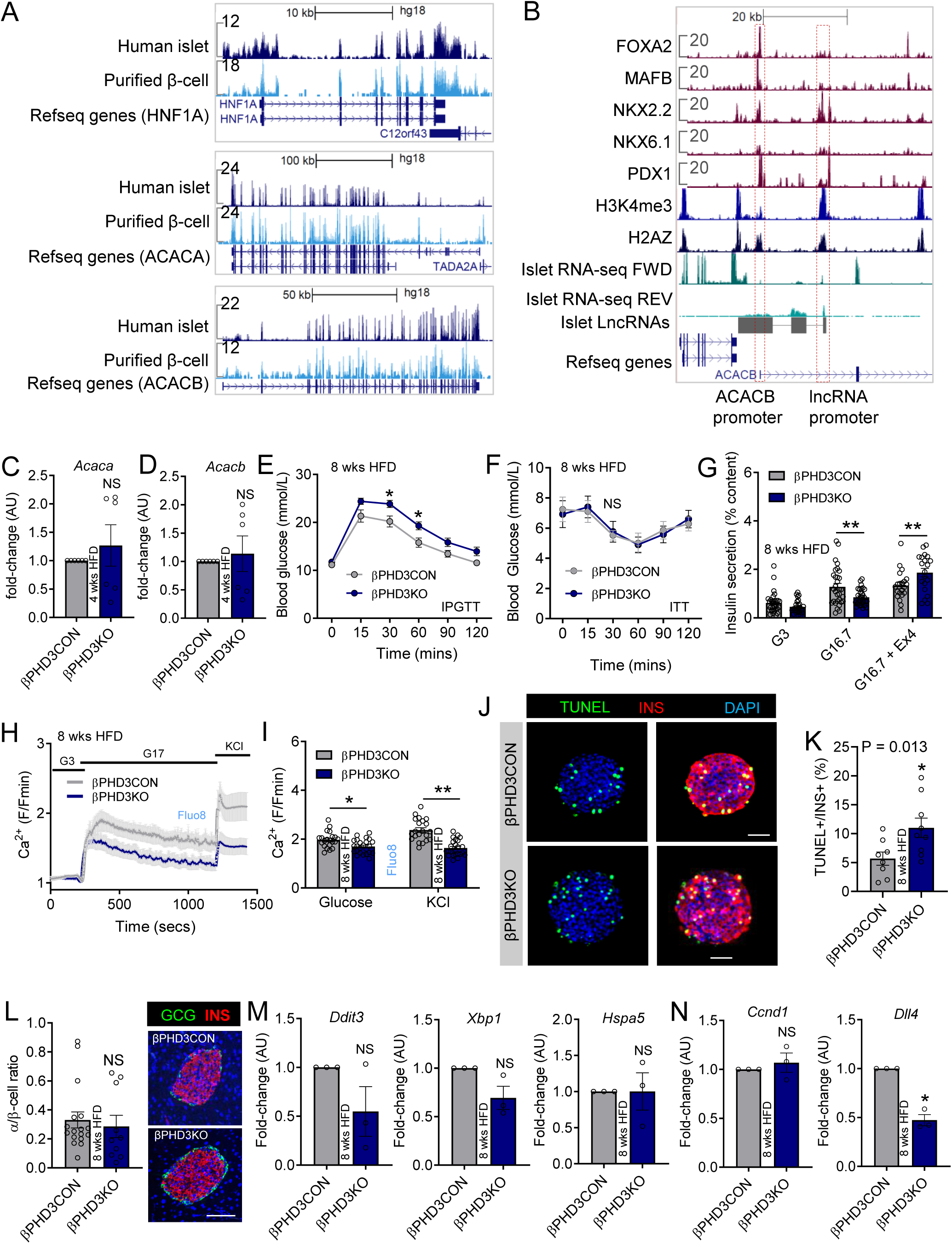
Prolonged metabolic stress induces failure in βPHD3KO islets. (A) *ACACB* is expressed in human islets and purified β-cells but levels are lower than *ACACA* (data were obtained from (35, 47, 48)). (B) The *ACACB* promoter is regulated by multiple β-cell transcription factors, with the presence of an antisense transcribed long non-coding RNA (detected data were also obtained from (35, 47, 48)). (C and D) Expression levels of *Acaca* (C) and *Acacb* (D) are not significantly different in βPHD3KO versus βPHD3CON islets following HFD (n = 6 animals) (paired t-test). (E) Glucose tolerance in βPHD3KO mice is still impaired versus βPHD3CON littermates after 8 weeks HFD (n = 9-11 animals/genotype) (two-way repeated measures ANOVA; Sidak’s multiple comparison test). (F) Insulin sensitivity is similar in 8 weeks HFD βPHD3CON and βPHD3KO mice (n = 5 animals/genotype) (two-way repeated measures ANOVA; Sidak’s multiple comparison test). (G) By 8 weeks of HFD, glucose-stimulated insulin secretion is impaired in βPHD3KO versus βPHD3CON islets. Note that Exendin-4 rescues the apparent secretory defect in βPHD3KO islets (n = 29-32 replicates from 4 animals/genotype) (two-way ANOVA, Bonferroni’s multiple comparison test). (H and I) Ca^2+^ responses to both glucose and KCl are impaired in 8 weeks HFD βPHD3KO islets, as shown by mean traces (H), as well as the summary bar graph (I) (n = 21-24 islets from 2 animals/genotype) (two-way ANOVA; Sidak’s multiple comparison test). (J and K) The proportion of apoptotic β-cells is increased in 8 weeks HFD βPHD3KO islets, shown by representative images (J) and summary bar graph (K) (scale bar = 42.5 µm) (n = 8-9 islets) (unpaired t-test). (L) α-cell/β-cell ratio is unchanged between 8 weeks HFD βPHD3CON and PHD3KO islets (scale bar = 42.5 µm) (n = 10-17 islets, 2 animals/genotype) (unpaired t-test). (M) Expression of the ER stress markers *Ddit3, Xbp1* and *Hspa5* is similar in 8 weeks HFD βPHD3CON and βPHD3KO islets (n = 3 animals/genotype) (paired t-test). (N) Expression of the HIF2α-targets *Ccnd1* and *Dll4* is unchanged or downregulated, respectively, in 8 weeks HFD βPHD3KO islets (n = 3 animals/genotype) (paired t-test). Bar graphs show scatter plot with mean ± SEM. Line graphs show mean ± SEM. *P<0.05, **P<0.01 and NS, non-significant. PHD3, prolyl-hydroxylase 3; Exendin-4, Ex4; G3, 3 mM glucose; G16.7, 16.7 mM glucose; G17, 17 mM glucose.

Closer examination of the promoter of *ACACB* gene in islets also revealed regulation by a number of established β-cell transcription factors, such as PDX1, MAFB and NKX2-2 (Figure 7B), further confirming the regulated expression of this gene. Unusually, a long non-coding RNA (lncRNA), transcribed antisense to the *ACACB* gene, was detected (Figure 7B), consistent with the presence of a negative regulatory mechanism for the expression of this gene in one or more cell types in the islet.

Suggesting that any regulation of ACC1/ACC2 by PHD3 is likely to be post-translational, as expected for a hydroxylase, qPCR analyses showed that both *Acaca* and *Acacb* expression were similar in 4 weeks HFD βPHD3CON and βPHD3KO islets (Figure 7C and D).

Thus, according to next generation sequencing, *ACACB* is reproducibly expressed in β-cells, but at levels lower than *ACACA*. Assuming that protein translation occurs, ACC2 might conceivably contribute to fatty acid oxidation in the absence of PHD3.

### PHD3 protects against lipotoxicity following prolonged metabolic stress

Lastly, we sought to investigate whether islets would eventually decompensate when faced with continued fatty acid/nutrient abundance. Glucose intolerance was still present in βPHD3KO mice following 8 weeks on HFD (Figure 7E), despite normal insulin sensitivity (Figure 7F). By this point, however, impaired glucose-stimulated insulin secretion (Figure 7G) was apparent in isolated βPHD3KO islets. This secretory deficit could be rescued by application of Ex4 to sensitize insulin granules to exocytosis (Figure 7G). In addition, the amplitude of glucose-stimulated Ca^2+^ rises was significantly reduced in 8 weeks HFD βPHD3KO compared to βPHD3CON islets (Figure 7H and I). Suggesting that profound defects in voltage-dependent Ca^2+^ channels might also be present, responses to the generic depolarizing stimulus KCl were markedly blunted in the same islets (Figure 7H and I). While apoptosis was increased in 8 weeks HFD βPHD3KO islets(Figure 7J and K), α-cell/β-cell ratio (Figure 7L) and expression of the ER stress markers *Ddit3, Xbp1* and *Hspa5* (Figure 7M) remained unchanged. Lastly, the HIF2α-target genes *Ccnd1* and *Dll4* were found to be either unchanged or downregulated in 8 weeks HFD βPHD3KO islets (Figure 7N), suggesting that HIF2α stabilization was unlikely to be the sole determinant of phenotype.

## DISCUSSION

In the present study, we build upon previous observations that chemical inhibition of all three PHD enzymes in islets and β-cell lines leads to alterations in glucose-stimulated insulin secretion (27). Specifically, we show that the alpha-ketoglutarate-dependent PHD3 maintains β-cell glucose sensing under states of metabolic stress and insulin resistance associated with fatty acid abundance. Our data suggest that PHD3 is required for ensuring that acetyl-CoA derived from glycolysis preferentially feeds the TCA cycle, linking blood glucose levels with ATP/ADP generation, β-cell electrical activity and insulin secretion. Loss of PHD3 leads to metabolic remodeling under HFD, resulting in a decrease in glycolytic fluxes, an increase in lactate accumulation and utilization of fatty acids as an energy source. This switch cannot be maintained, however, and β-cells eventually fail following prolonged exposure to fatty acids. Thus, PHD3 appears to be a critical component of the β-cell metabolic machinery required for glucose sensing during episodes of nutritional overload.

How does PHD3 maintain glucose metabolism in β-cells? Previous studies in cancer cells have shown that PHD3 hydroxylates and activates ACC2, suppressing beta oxidation (24). While β-cells are thought to predominantly express ACC1, the levels of *ACACB*, which encodes ACC2, were found to be similar to the β-cell transcription factor HNF1A, albeit much lower than for *ACACA*. Intriguingly, *ACACB* was enriched for promoter sites suggestive of negative regulation, which is unusual amongst β-cell genes, and *Acaca/Acacb* were not upregulated in HFD βPHD3CON or βPHD3KO islets. This supports a potential role for post-transcriptional modifications in determining ACC2 activity. We thus propose that loss of PHD3 might lead to suppression of ACC2 activity, which becomes apparent during HFD when its substrate is present in abundance. Alternatively, PHD3 might hydroxylate and activate ACC1, leading to regulation of CPT1 by malonyl-CoA when fatty acids are supplied in excess, as suggested by glucose oxidation experiments. In any case, experiments with etomoxir strongly infer a role for CPT1 in the negative effects of PHD3 deletion on glucose metabolism. While etomoxir has been shown to target complex I of the electron transport to lower ATP production (49), we don’t think this played a major role here, since ATP levels were restored in βPHD3KO islets treated with the inhibitor.

dentifying the efficiency of protein translation for *ACACB* → ACC2, as well as the PHD3 hydroxylation sites involved, will be critical. However, currently available antibodies for detection of ACC2, as well its hydroxylated forms, are poor. Moreover, assigning hydroxylation targets using mass spectrometry is controversial: mis-alignment of hydroxylation is commonly associated with the presence of residues in the tryptic fragment that can be artefactually oxidised (50). Thus, studies using animals lacking both PHD3 and ACC2 specifically in β-cells would be required to definitively link the carboxylase with the phenotype here.

As normal chow contains a low proportion of calories from fat, metabolic stress was needed to reveal the full *in vitro* and *in vivo* phenotype associated with PHD3 loss. These data also support an effect of PHD3 on ACC1/2 and CPT1, since without acyl-CoA derived from exogenous fatty acids, glucose would still constitute the primary fuel source and regulator of insulin release. The lack of phenotype under normal diet is unlikely to reflect the age of the animals, since even at 20 weeks of age, glucose intolerance was still not present in βPHD3KO mice. An alternative explanation is that loss of PHD3 can be compensated under normal conditions, while other mechanisms associated with fatty acid excess and lipogenesis, for example ER stress (51, 52), also contribute to the βPHD3KO phenotype. We feel that this explanation is less likely, however, since we could not detect upregulation of *Ddit3, Xbp1* and *Hspa5* even following 8 weeks HFD.

Suggesting that the phenotype associated with PHD3 loss was not solely due to HIF, no differences in the gene expression of HIF1 targets could be detected in βPHD3KO versus βPHD3CON islets. Indeed, PHD2 is the major hydroxylase that regulates HIF1α stability (11, 12), with no changes in activity of the transcription factor detected following PHD3 loss (11, 12, 53). Thus, it is perhaps unsurprising that there is a lack of HIF1 transcriptional signature in βPHD3KO islets, in agreement with previous studies in other tissues (53, 54). In addition, glucose-stimulated Ca^2+^ fluxes, a sensitive readout of changes in oxygen-dependent regulation (38), were unaffected during hypoxia in βPHD3KO islets. Lastly, no changes in expression of the HIF1-sensitive gene *Ldha* (55) were detected. Nonetheless, we cannot completely exclude HIF-dependent effects, and as such, studies should either be repeated on a HIF1- and HIF2-null background (i.e. a quadruple transgenic) or using (moderately) specific chemical inhibitors.

An intriguing observation was that PHD3 deletion decreased Exendin-4-but not glucose-stimulated insulin secretion in islets from animals fed standard chow. Given that *Glp1r* expression and signaling remained intact in βPHD3KO islets, alterations in cytosolic glutamate accumulation might instead be present, previously shown to prime incretin-responsiveness following its release with insulin from the granule (56). Arguing against this, however, interrogation of the metabolic tracing data showed that steady-state glutamate levels were unchanged between βPHD3CON and βPHD3KO islets, meaning that glucose was still able to enter the malate-aspartate shuttle to produce the neurotransmitter. It will be interesting in the future to pinpoint how PHD3 impinges upon Ex4-potentiated insulin secretion.

In summary, PHD3 possesses a conserved role in gating nutrient preference toward glucose and glycolysis during both cell transformation (24) and metabolic stress (here). It will be interesting to now study whether similar effects of PHD3 are present in other cell types involved in glucose-sensing (for example, pancreatic alpha cells, hypothalamic neurons).

## MATERIALS AND METHODS

### Study design

No data were excluded unless the cells displayed a non-physiological state (i.e. impaired viability). All individual data points are reported. The measurement unit is animal or batch of islets, with experiments replicated independently. Animals and islets were randomly allocated to treatment groups to ensure that all states were represented in the different experiment arms.

### Study approval

Animal studies were regulated by the Animals (Scientific Procedures) Act 1986 of the U.K., and approval was granted by the University of Birmingham’s Animal Welfare and Ethical Review Body.

### Mouse models

β-cell-specific PHD3 (βPHD3KO) knockout mice were generated using the Cre-LoxP system. *Ins1Cre* mice (JAX stock no. 026801), with Cre-recombinase knocked into the *Ins1* gene locus (34), were crossbred to mice carrying *flox’d* alleles for PHD3 (*Egln3*^*fl/fl*^) (9). Adult male and female βPHD3KO animals (*Ins1Cre*^*+/-*^;*Egln3*^*fl/fl*^) and their controls (βPHD3CON) (*Ins1*^*wt/wt-*^; *Egln3*^*fl/fl*^, *Ins1Cre*^*+/-*^;*Egln3*^*wt/wt*^ and *Ins1*^*wt/wt*^;*Egln3*^*wt/wt*^) were used from 8-20 weeks of age. No extra-pancreatic recombination has been observed in *Ins1Cre* mice and possession of a *Cre* allele is not associated with any changes in glucose homeostasis in our hands (33, 57). Given recently reported issues with allele-silencing in some *Ins1Cre* colonies (58), recombination efficiency of our line was regularly monitored and verified to be >90% using ROSA^mT/mG^ reporter animals (59). Animals were maintained on a C57BL/6J background and backcrossed for at least 6 generations following re-derivation into the animal facility. Lines were regularly refreshed by crossing to bought-in C57BL/6J (Charles River). βPHD3CON and βPHD3KO mice were fed standard chow (SC) and/or high fat diet containing 60% fat (HFD), with body weight checked weekly until 18 weeks of age. Animals were maintained in a specific pathogen-free facility, with free access to food and water.

### Intraperitoneal and oral glucose tolerance testing

Mice were fasted for 4-6 hours, before intraperitoneal injection of glucose (1-2 g/kg body weight). Blood samples for glucose measurement were taken from the tail vein at 0, 15, 30, 60, 90 and 120 min post-challenge. Glucose was measured using a Contour XT glucometer (Bayer, Germany). For mice on SC, intraperitoneal glucose tolerance testing (IPGTT) was performed every 2-4 weeks, between 8-20 weeks of age. HFD-fed mice underwent IPGTT following 4 and 8 weeks of HFD. Oral glucose tolerance testing (OGTT) was performed as for IPGTT, except that 2 g/kg body weight glucose was delivered using an oral gavage tube.

### Serum insulin

Blood samples were collected following intraperitoneal glucose injection (1-2 g/kg body weight). Serum was separated by centrifugation (2500 rpm/10 min/4°C), before assaying using a HTRF Mouse Serum Insulin Assay kit assay (Cisbio, France). Due to NC3R limits on blood sample volumes, insulin was only measured at 0, 15 and 30 min post-glucose injection.

### Insulin tolerance test (ITT)

Mice fasted for 4-6 hours (SC cohort) or 14-16 hours (HFD cohort) received 0.75 U/kg body weight insulin (Humulin S, 100 U/ml, Lilly, UK) given by intraperitoneal injection. Blood glucose was measured at 0, 15, 30, 60, 90 and 120 min post-insulin injection.

### Islet isolation

Islets were isolated following bile duct injection with Serva NB8 1 mg/ml collagenase and Histopaque/Ficoll gradient separation. Islets were cultured in RPMI medium containing 10% FCS, 100 units/mL penicillin, and 100 μg/mL streptomycin at 5% CO_2_, 37°C. For experiments under hypoxia, islets were incubated in a Don Whitely H35 Hypoxystation, allowing oxygen tension to be finely regulated at either 1% or 21%. For experiments with exogenous lipids, islets were treated with either 0.75% bovine serum albumin (BSA) control, or 150µM sodium palmitate dissolved in 0.75% BSA.

### Gene expression

Trizol purification was used for mRNA extraction, while cDNA was synthesized by reverse transcription. Gene expression was detected by real time PCR (qPCR), using PowerUp SYBR Green Master Mix (Thermofisher Scientific) and quantification was based on the 2^−ΔΔCt^ method, expressed as fold-change in gene expression. The following primers were used: *Ppia* (forward 5’ AAGACTGAGTGGTTGGATGG 3’, reverse 5’ ATGGTGATCTTCTTGCTGGT 3’), *Actb* (forward 5’ CGAGTCGCGTCCACCC 3’, reverse 5’ CATCCATGGCGAACTGGTG 3’), *Egln3* (beginning of exon2) (forward 5’ GCTTGCTATCCAGGAAATGG 3’, reverse 5’ GCGTCCCAATTCTTATTCAG 3’), *Egln3* (end of exon1) (forward 5’ GGCTGGGCAAATACTATGTCAA 3’, reverse 5’ GGTTGTCCACATGGCGAACA 3’), *Bnip3* (forward 5’ CTGGGTAGAACTGCACTTCAG 3’, reverse 5’ GGTTGTCCACATGGCGAACA 3’), *Car9* (forward 5’ GGAGCTACTTCGTCCAGATTCAT 3’, reverse 5’ CCGGAACTGAGCCTATCCAAC 3’), *Gls* (forward 5’ TTCGCCCTCGGAGATCCTAC 3’, reverse 5’ CCAAGCTAGGTAACAGACCCT 3’), *Ldha* (forward 5’ TTCAGCGCGGTTCCGTTAC 3’, reverse 5’ CCGGCAACATTCACACCAC 3’), *Cpt1a* (forward 5’ CTCCGCCTGAGCCATGAAG 3’, reverse 5’ CACCAGTGATGATGCCATTCT 3’), *Acaca* (forward 5’ CTTCCTGACAAACGAGTCTGG 3’, reverse 5’ CTGCCGAAACATCTCTGGGA 3’), *Acacb* (forward 5’ CCTTTGGCAACAAGCAAGGTA 3’, reverse 5’ AGTCGTACACATAGGTGGTCC 3’), *Pdx1* (forward 5’ CCAAAGCTCACGCGTGGA 3’, reverse 5’ TGTTTTCCTCGGGTTCCG 3’), *Nkx6-1* (forward 5’ GCCTGTACCCCCCATCAAG 3’, reverse 5’ GTGGGTCTGGTGTGTTTTCTCTT 3’), *Mafa* (forward 5’ CTTCAGCAAGGAGGAGGTCATC 3’, reverse 5’ CGTAGCCGCGGTTCTTGA 3’), *Ddit3* (forward 5’ CTGGAAGCCTGGTATGAGGAT 3’, reverse 5’ CAGGGTCAAGAGTAGTGAAGGT 3’), *Xbp1* (forward 5’ AGCAGCAAGTGGTGGATTTG 3’, reverse 5’ GAGTTTTCTCCCGTAAAAGCTGA 3’), *Hspa5*, (forward 5’ ACTTGGGGACCACCTATTCCT 3’, reverse 5’ GTTGCCCTGATCGTTGGCTA 3’), *Ccnd1*, (forward 5’ GCGTACCCTGACACCAATCTC 3’, reverse 5’ CTCCTCTTCGCACTTCTGCTC 3’) and *Dll4*, (forward 5’ TTCCAGGCAACCTTCTCCGA 3’, reverse 5’ ACTGCCGCTATTCTTGTCCC 3’).

### Western Blot

Proteins of interest were lysed in 1x Laemmli buffer and separated using SDS-PAGE before transfer onto a nitrocellulose membrane. After blocking, primary rabbit anti-PHD3 1:1000 (Abcam, ab184714) or mouse anti-actin 1:2000 (Sigma-Aldrich, A4700) antibodies were diluted in 5% milk before incubation overnight on a shaker. Detection was performed using secondary goat anti-rabbit IgG 1:1000 (Cell Signaling, 7074S) or horse anti-mouse IgG 1:2000 (Cell Signaling, 7076S) and ECL. Protein quantification was based on blot intensity, measured using ImageJ (NIH).

### Immunohistochemistry

Pancreata were isolated, fixed in 10% formalin and embedded in paraffin. Paraffin slides were deparaffinized and rehydrated, before antigen retrieval using citrate buffer. Sections were incubated overnight at 4°C with rabbit anti-insulin 1:500 (Cell Signaling, 3014S) and mouse anti-glucagon 1:2000 (Sigma-Aldrich, G 2654) followed by washing and application of goat anti-rabbit Alexa Fluor 647 1:500 (ThermoFisher Scientific, A-21244) and goat anti mouse DyLight 488 1:500 (Invitrogen, 35503). Coverslips were mounted using VECTASHIELD HardSet with Dapi and 425 images per section captured using a Zeiss Axio Scan.Z1 automated slide scanner equipped with a 20 x / 0.8 NA objective. β-cell mass (%) was calculated as the area of insulin + staining/area of the pancreas. Excitation was delivered at λ = 330-375 nm and λ = 590-650 nm for DAPI and Alexa Fluor 647, respectively. Emitted signals were detected using an Orca Flash 4.0 at λ = 430-470 nm and λ = 663-738 nm for DAPI and Alexa Fluor 647, respectively.

TUNEL staining was performed using the DeadEnd Fluorometric TUNEL System (Promega), as previously described (60). The proportion of apoptotic β-cells was calculated as the area of TUNEL+ staining (fluorescein-12-dUTP)/area of insulin+ staining (as above). α-cell/ β-cell ratio was calculated following staining with antibodies against insulin (as above) and glucagon (primary antibody: mouse anti-glucagon 1:2000; Sigma-Aldrich, G2645) (secondary antibody goat anti-mouse Alexa Fluor 488 1:500; ThermoFisher Scientific, A11001). Images were captured using a Zeiss LSM780 meta-confocal microscope equipped with highly-sensitive GaAsP PMT detectors. Excitation was delivered at λ = 405 nm, λ = 488 nm and λ = 633 nm for DAPI, fluorescein-12-dUTP/Alexa Fluor 488 and Alexa Fluor 647 nm, respectively. Emitted signals were detected at λ = 428-533 nm, λ = 498-559 nm and λ = 643–735 nm for DAPI, fluorescein-12-dUTP/Alexa Fluor 488 and Alexa Fluor 633 nm, respectively.

### Insulin secretion *in vitro* and insulin measurement

Ten to fifteen size-matched islets were stimulated with: 3 mM glucose, 16.7 mM glucose and 16.7 mM glucose + 10 nM Exendin-4 in HEPES-bicarbonate buffer (mM: 120 NaCl, 4.8 KCl, 24 NaHCO_3_, 0.5 Na_2_HPO_4_, 5 HEPES, 2.5 CaCl_2_, 1.2 MgCl_2_) supplemented with 0.1% BSA at 37°C. Insulin content was extracted using acid ethanol. Insulin concentration (ng/ml) was measured using a HTRF Insulin Ultra-Sensitive Assay kit (Cisbio, France).

### Live imaging

Islets were loaded with the Ca^2+^ indicators Fluo8 or Fura2 (both AAT Bioquest), before imaging using a Crest X-Light spinning disk microscope coupled to a Nikon Ti-E base with 10 x 0.4 NA and 20 x 0.8 NA objectives. For Fluo8 imaging, excitation was delivered at and λ = 458–482 nm using a Lumencor Spectra X light engine. Emission was captured at λ = 500-550 nm using a highly-sensitive Photometrics Delta Evolve EM-CCD. For experiments with the ratiometric Ca^2+^ indicator, Fura2, excitation was delivered at λ = 340 nm and λ = 385 nm using Cairn Research Fura LEDs in widefield mode, with emitted signals detected at λ = 470–550 nm.

For ATP/ADP imaging, islets were transduced with the ATP/ADP sensor Perceval (a kind gift from Prof Gary Yellen, Harvard) (61) using an adenoviral vector and imaged identically to Fluo8. For FRET-based cAMP imaging, islets were infected with adenovirus harboring Epac2-camps (a kind gift from Prof Dermot Cooper, Cambridge). Excitation was delivered at 430– 450 nm, with emission detected at and λ = 460–500 and and λ = 520–550 nm for Cerulean and Citrine, respectively.

In all cases, HEPES-bicarbonate buffer was used (mM: 120 NaCl, 4.8 KCl, 24 NaHCO_3_, 0.5 Na_2_HPO_4_, 5 HEPES, 2.5 CaCl_2_, 1.2 MgCl_2_, and 3–17 D-glucose), with glucose and drugs being added at the indicated concentrations and timepoints. Fura2 and Epac2-camps traces were normalized as the ratio of 340/385 or Cerulean/Citrine, respectively. Data were presented as raw or F/F_min_ where F = fluorescence at any timepoint and F_min_ = minimum fluorescence, or R/R_0_ where R = fluorescence at any timepoint and R_0_ = fluorescence at 0 mins.

### Glucose oxidation assays and metabolic tracing

^14^C glucose oxidation and lipid incorporation: batches of 40 islets were used for quantification of glucose oxidation and incorporation into lipids by scintillation spectrometry, as previously described (44).

Gas chromatography–mass spectrometry (GC-MS)-based ^13^C_6_ mass isotopomer distribution: To ensure steady state, 50-100 islets were cultured with 10 mM ^13^C_6_-glucose for 24 hours (62), before extraction of metabolites using sequentially pre-chilled HPLC-grade methanol, HPLC-grade distilled H_2_O containing 1 μg/mL D6-glutaric acid and HPLC-grade chloroform at −20 °C. Polar fractions were separated by centrifugation, vacuum dried and solubilized in 2% methoxyamine hydrochloric acid in pyridine. Samples were derivatized using N-tertbutyldimethylsilyl-N-methyltrifluoroacetamide (MTBSTFA) with 1% (w/v) tertbutyldimethyl-chlorosilane (TBDMCS), before analysis on an Agilent 7890B gas chromatograph mass spectrometer, equipped with a medium polar range polydimethylsiloxane GC column (DB35-MS). Mass isotopomer distributions (MIDs) were determined based upon spectra corrected for natural isotope abundance. Data were analyzed using MetaboliteDetector software (63).

### Statistics

Measurements were performed on discrete samples unless otherwise stated. Data normality was assessed using D’Agostino-Person test. All analyses were conducted using GraphPad Prism software. Pairwise comparisons were made using Student’s unpaired or paired t-test, or Mann-Whitney test. Multiple interactions were determined using either Kruskal-Wallis test, one-way ANOVA or two-way ANOVA followed by Tukey’s, Dunn’s, Dunnett’s, Bonferonni’s or Sidak’s post-hoc tests (accounting for degrees of freedom).

### Data availability

The datasets generated and/or analyzed during the current study are available from the corresponding author upon reasonable request.

## ACKNOWLEDGEMENTS

D.J.H. was supported by a Diabetes UK R.D. Lawrence (12/0004431) Fellowship, a Wellcome Trust Institutional Support Award, and MRC (MR/N00275X/1 and MR/S025618/1) and Diabetes UK (17/0005681) Project Grants. D.T. was supported by Cancer Research UK Grants (C42109/A26982 and C42109/A24891). This project has received funding from the European Research Council (ERC) under the European Union’s Horizon 2020 research and innovation programme (Starting Grant 715884 to D.J.H.). We thank Dr Mathew Coleman (University of Birmingham) for useful discussions.

## AUTHOR CONTRIBUTIONS

D.N., F.C., R.B.B., R.W., and J.C. performed experiments and analyzed data. F.C. and A.T. ran and analyzed samples for GC-MS. I.A. analyzed data. D.T. and D.J.H. conceived, designed and supervised the studies. D.J.H. and D.T. wrote the paper with input from all authors.

## REFERENCES

1. R. K. Bruick, Oxygen sensing in the hypoxic response pathway: regulation of the hypoxia-inducible transcription factor. Genes Dev. 17, 2614–2623 (2003).

2. C. J. Schofield, P. J. Ratcliffe, Oxygen sensing by HIF hydroxylases. Nat. Rev. Mol. Cell Biol. 5, 343–354 (2004).

3. R. K. Bruick, S. L. McKnight, A conserved family of prolyl-4-hydroxylases that modify HIF. Science 294, 1337–1340 (2001).

4. A. C. Epstein et al., C. elegans EGL-9 and mammalian homologs define a family of dioxygenases that regulate HIF by prolyl hydroxylation. Cell 107, 43–54 (2001).

5. W. G. Kaelin, Jr., P. J. Ratcliffe, Oxygen sensing by metazoans: the central role of the HIF hydroxylase pathway. Mol. Cell 30, 393–402 (2008).

6. D. A. Chan et al., Tumor vasculature is regulated by PHD2-mediated angiogenesis and bone marrow-derived cell recruitment. Cancer Cell 15, 527–538 (2009).

7. R. V. Duran et al., HIF-independent role of prolyl hydroxylases in the cellular response to amino acids. Oncogene 32, 4549–4556 (2013).

8. W. Luo et al., Pyruvate kinase M2 is a PHD3-stimulated coactivator for hypoxia-inducible factor 1. Cell 145, 732–744 (2011).

9. A. T. Henze et al., Loss of PHD3 allows tumours to overcome hypoxic growth inhibition and sustain proliferation through EGFR. Nat. Commun. 5, 5582 (2014).

10. R. J. Appelhoff et al., Differential Function of the Prolyl Hydroxylases PHD1, PHD2, and PHD3 in the Regulation of Hypoxia-inducible Factor. J. Biol. Chem. 279, 38458–38465 (2004).

11. E. Berra et al., HIF prolyl-hydroxylase 2 is the key oxygen sensor setting low steady-state levels of HIF-1alpha in normoxia. EMBO J 22, 4082–4090 (2003).

12. D. A. Tennant et al., Reactivating HIF prolyl hydroxylases under hypoxia results in metabolic catastrophe and cell death. Oncogene 28, 4009–4021 (2009).

13. J. Kiss et al., Loss of the oxygen sensor PHD3 enhances the innate immune response to abdominal sepsis. J. Immunol. 189, 1955–1965 (2012).

14. S. R. Walmsley et al., Prolyl hydroxylase 3 (PHD3) is essential for hypoxic regulation of neutrophilic inflammation in humans and mice. J. Clin. Invest. 121, 1053–1063 (2011).

15. Y. Su et al., PHD3 regulates differentiation, tumour growth and angiogenesis in pancreatic cancer. Br. J. Cancer 103, 1571–1579 (2010).

16. S. Schlisio et al., The kinesin KIF1Bbeta acts downstream from EglN3 to induce apoptosis and is a potential 1p36 tumor suppressor. Genes Dev. 22, 884–893 (2008).

17. T. L. Place, F. E. Domann, Prolyl-hydroxylase 3: Evolving Roles for an Ancient Signaling Protein. Hypoxia (Auckl) 2013, 13–17 (2013).

18. D. A. Tennant, E. Gottlieb, HIF prolyl hydroxylase-3 mediates alpha-ketoglutarate-induced apoptosis and tumor suppression. J. Mol. Med. 88, 839–849 (2010).

19. H. Boulahbel, Raúl V. Durán, E. Gottlieb, Prolyl hydroxylases as regulators of cell metabolism: Figure 1. Biochem. Soc. Trans. 37, 291–294 (2009).

20. N. Chen et al., The oxygen sensor PHD3 limits glycolysis under hypoxia via direct binding to pyruvate kinase. Cell Res. 21, 983–986 (2011).

21. S. Lee et al., Neuronal apoptosis linked to EglN3 prolyl hydroxylase and familial pheochromocytoma genes: Developmental culling and cancer. Cancer Cell 8, 155–167 (2005).

22. L. Dang et al., Cancer-associated IDH1 mutations produce 2-hydroxyglutarate. Nature 462, 739–744 (2009).

23. I. P. Tomlinson et al., Germline mutations in FH predispose to dominantly inherited uterine fibroids, skin leiomyomata and papillary renal cell cancer. Nat Genet 30, 406–410 (2002).

24. N. J. German et al., PHD3 Loss in Cancer Enables Metabolic Reliance on Fatty Acid Oxidation via Deactivation of ACC2. Mol. Cell 63, 1006–1020 (2016).

25. C. M. Taniguchi et al., Cross-talk between hypoxia and insulin signaling through Phd3 regulates hepatic glucose and lipid metabolism and ameliorates diabetes. Nat. Med. 19, 1325–1330 (2013).

26. H. Yano et al., PHD3 regulates glucose metabolism by suppressing stress-induced signalling and optimising gluconeogenesis and insulin signalling in hepatocytes. Scientific Reports 8 (2018).

27. M. Huang et al., Role of prolyl hydroxylase domain proteins in the regulation of insulin secretion. Physiological Reports 4, e12722 (2016).

28. A. De Vos et al., Human and rat beta cells differ in glucose transporter but not in glucokinase gene expression. J. Clin. Invest. 96, 2489–2495 (1995).

29. M. S. German, Glucose sensing in pancreatic islet beta cells: the key role of glucokinase and the glycolytic intermediates. Proc. Natl. Acad. Sci. U. S. A. 90, 1781–1785 (1993).

30. G. A. Rutter, T. J. Pullen, D. J. Hodson, A. Martinez-Sanchez, Pancreatic beta-cell identity, glucose sensing and the control of insulin secretion. Biochem. J. 466, 203–218 (2015).

31. T. J. Pullen, G. A. Rutter, When less is more: the forbidden fruits of gene repression in the adult beta-cell. Diabetes Obes. Metab. 15, 503–512 (2013).

32. K. Lemaire, L. Thorrez, F. Schuit, Disallowed and Allowed Gene Expression: Two Faces of Mature Islet Beta Cells. Annu. Rev. Nutr. 36, 45–71 (2016).

33. B. Thorens et al., Ins1 knock-in mice for beta cell-specific gene recombination. Diabetologia 58, 558–565 (2014).

34. K. Takeda et al., Placental but Not Heart Defects Are Associated with Elevated Hypoxia-Inducible Factor α Levels in Mice Lacking Prolyl Hydroxylase Domain Protein 2. Mol. Cell. Biol. 26, 8336–8346 (2006).

35. D. M. Blodgett et al., Novel Observations From Next-Generation RNA Sequencing of Highly Purified Human Adult and Fetal Islet Cell Subsets. Diabetes 64, 3172–3181 (2015).

36. C. Benner et al., The transcriptional landscape of mouse beta cells compared to human beta cells reveals notable species differences in long non-coding RNA and protein-coding gene expression. BMC Genomics 15 (2014).

37. H. Komatsu et al., Oxygen environment and islet size are the primary limiting factors of isolated pancreatic islet survival. PLoS ONE 12, e0183780 (2017).

38. J. Cantley et al., Deletion of the von Hippel–Lindau gene in pancreatic β cells impairs glucose homeostasis in mice. J. Clin. Invest. 10.1172/jci26934 (2008).

39. M. A. Nauck et al., Incretin effects of increasing glucose loads in man calculated from venous insulin and C-peptide responses. J. Clin. Endocrinol. Metab. 63, 492–498 (1986).

40. J. Cantley, S. T. Grey, P. H. Maxwell, D. J. Withers, The hypoxia response pathway and β-cell function. Diabetes Obes. Metab. 12, 159–167 (2010).

41. F. Dayan, D. Roux, M. C. Brahimi-Horn, J. Pouyssegur, N. M. Mazure, The oxygen sensor factor-inhibiting hypoxia-inducible factor-1 controls expression of distinct genes through the bifunctional transcriptional character of hypoxia-inducible factor-1alpha. Cancer Res 66, 3688–3698 (2006).

42. G. da Silva Xavier, D. J. Hodson, Mouse models of peripheral metabolic disease. Best Practice & Research Clinical Endocrinology & Metabolism 10.1016/j.beem.2018.03.009 (2018).

43. N. M. Templeman et al., Reduced Circulating Insulin Enhances Insulin Sensitivity in Old Mice and Extends Lifespan. Cell Reports 20, 451–463 (2017).

44. J. Cantley et al., Disruption of beta cell acetyl-CoA carboxylase-1 in mice impairs insulin secretion and beta cell mass. Diabetologia 62, 99–111 (2018).

45. S. M. Ronnebaum et al., Chronic Suppression of Acetyl-CoA Carboxylase 1 in β-Cells Impairs Insulin Secretion via Inhibition of Glucose Rather Than Lipid Metabolism. J. Biol. Chem. 283, 14248–14256 (2008).

46. M. J. MacDonald, A. Dobrzyn, J. Ntambi, S. W. Stoker, The role of rapid lipogenesis in insulin secretion: Insulin secretagogues acutely alter lipid composition of INS-1 832/13 cells. Arch. Biochem. Biophys. 470, 153–162 (2008).

47. S. Hrvatin et al., Differentiated human stem cells resemble fetal, not adult, β cells. Proceedings of the National Academy of Sciences 111, 3038–3043 (2014).

48. I. Moran et al., Human beta Cell Transcriptome Analysis Uncovers lncRNAs That Are Tissue-Specific, Dynamically Regulated, and Abnormally Expressed in Type 2 Diabetes. Cell Metab. 16, 435–448 (2012).

49. B. Raud et al., Etomoxir Actions on Regulatory and Memory T Cells Are Independent of Cpt1a-Mediated Fatty Acid Oxidation. Cell Metab. 28, 504-515.e507 (2018).

50. M. E. Cockman et al., Lack of activity of recombinant HIF prolyl hydroxylases (PHDs) on reported non-HIF substrates. eLife 8 (2019).

51. M. Cnop et al., RNA Sequencing Identifies Dysregulation of the Human Pancreatic Islet Transcriptome by the Saturated Fatty Acid Palmitate. Diabetes 63, 1978–1993 (2014).

52. D. A. Cunha et al., Initiation and execution of lipotoxic ER stress in pancreatic beta-cells. J. Cell Sci. 121, 2308–2318 (2008).

53. C. M. Taniguchi et al., Cross-talk between hypoxia and insulin signaling through Phd3 regulates hepatic glucose and lipid metabolism and ameliorates diabetes. Nat Med 19, 1325–1330 (2013).

54. T. Bishop et al., Abnormal sympathoadrenal development and systemic hypotension in PHD3-/-mice. Mol Cell Biol 28, 3386–3400 (2008).

55. X.-g. Cui et al., HIF1/2α mediates hypoxia-induced LDHA expression in human pancreatic cancer cells. Oncotarget 8 (2017).

56. G. Gheni et al., Glutamate Acts as a Key Signal Linking Glucose Metabolism to Incretin/cAMP Action to Amplify Insulin Secretion. Cell Reports 9, 661–673 (2014).

57. Natalie R. Johnston et al., Beta Cell Hubs Dictate Pancreatic Islet Responses to Glucose. Cell Metab. 24, 389–401 (2016).

58. M. L. Golson et al., Ins1-Cre and Ins1-CreER gene replacement alleles are susceptible to silencing by DNA hypermethylation. Endocrinology 10.1210/endocr/bqaa054 (2020).

59. J. Ast et al., Super-resolution microscopy compatible fluorescent probes reveal endogenous glucagon-like peptide-1 receptor distribution and dynamics. Nat. Commun. 11 (2020).

60. D. J. Hodson et al., ADCY5 couples glucose to insulin secretion in human islets. Diabetes 63, 3009–3021 (2014).

61. J. Berg, Y. P. Hung, G. Yellen, A genetically encoded fluorescent reporter of ATP:ADP ratio. Nat. Methods 6, 161–166 (2009).

62. M. Wortham et al., Integrated In Vivo Quantitative Proteomics and Nutrient Tracing Reveals Age-Related Metabolic Rewiring of Pancreatic β Cell Function. Cell Reports 25, 2904-2918.e2908 (2018).

63. K. Hiller et al., MetaboliteDetector: Comprehensive Analysis Tool for Targeted and Nontargeted GC/MS Based Metabolome Analysis. Anal. Chem. 81, 3429–3439 (2009).

